# Tradeoffs in the Design of RNA Thermometers

**DOI:** 10.1101/2023.11.29.568956

**Authors:** Krishan Kumar Gola, Abhilash Patel, Shaunak Sen

## Abstract

The synthesis of RNA thermometers is aimed at achieving temperature responses with desired thresholds and sensitivities. Although previous works have generated thermometers with a variety of thresholds and sensitivities as well as guidelines for design, possible constraints in the achievable thresholds and sensitivities remain unclear. We addressed this issue using a two-state model and its variants, as well as melt profiles generated from thermodynamic computations. In the two-state model, we found that the threshold was inversely proportional to the sensitivity, in the case of a fixed energy difference between the two states. Notably, this constraint could persist in variations of the two-state model with sequentially unfolding states and branched parallel pathways. Furthermore, the melt profiles generated from a library of thermometers exhibited a similar constraint. These results should inform the design of RNA thermometers as well as other responses that are mediated in a similar fashion.

## Introduction

RNA thermometers are RNA elements that change their conformation in a temperature-dependent fashion.^1^ Moreover, this conformational change triggers a downstream response. A typical example of this phenomenon is a temperature-dependent conformational change that enables the ribosome to access the ribosome binding site (RBS) and initiate translation. Furthermore, the change in the activity of an RNA thermometer is an important signal of the environmental temperature. An early report of an RNA thermometer was in the *λ cIII* gene.^2^ RNA thermometers have also been found to regulate responses associated with temperature change such as the heat-shock response, the cold-shock response and the virulence of pathogens.^3^ Hence, it is likely that the properties of these thermometers are tuned by cells to achieve the desired temperature response.

The threshold temperature at which the temperature response is triggered can vary. Usually, a shift to 37 °C, signifying the body temperature of the host mammal, is considered to be an important threshold for pathogenic bacteria.^1^ For example, the *prfA* thermometer has a different response at temperatures below 30 °C in comparison with temperatures above 37 °C.^4^ For other RNA thermometers, the RBS is accessible to the ribosome only at 42 °C.^5^ Furthermore, *Synechocystis* cells exhibit a large induction of *hsp17* mRNA when the temperature changes from 34 °C to 44 °C.^6^ The sensitivity of such responses has also been noted as an important functional property.^6^ For instance, *prfA* has been reported to exhibit a large change in response over a narrow temperature range, while *cssA* has been found to exhibit a relatively gradual response.^4^ The ability of RNA thermometers to detect changes as small as 1 °C has also been highlighted.^1^ In general, this feature can be characterized in terms of the sensitivity of the response, or the amount of change in the output for a small change in the input. In this context, co-operative melting transitions have often been noted as underlying the sensitivity in these responses.^7^ Moreover, different structural elements contributing to the temperature-sensing property of these thermometers have been identified. ^1^ Several synthetic designs have also been made,^8–12^ which are mostly simpler than naturally occurring thermometers and typically have smaller fold changes compared to similar elements made with proteins.^13^ In such cases, the thermometer responses have been observed to lead to a qualitative shift in sensitivities even with one base change in the thermometer sequences.^8,10^ A key design specification for such thermometers is to obtain a response of the desired sensitivity and threshold, which can be tuned in the relevant physiological range.^8^

There are at least three striking aspects with regard to the design space of RNA thermometers. The first aspect is the immense structural diversity underlying a temperature response, both in the minimum free energy structures of different thermometers and in the potentially large ensemble of structures for a single thermometer. The second aspect is the diverse temperatures at which they trigger a response, ranging at least from 30 °C to 44 °C. The third aspect is the possible functional significance of other dimensions in the performance space of the RNA thermometer responses, such as the sensitivity, or the extent of the change in the response to a specific change in the temperature. However, the existence of constraints in this design space, especially with regard to the different threshold temperatures and possible maximum sensitivities, is generally unclear.

Here we asked whether there were constraints to the co-variation of the RNA thermometer response properties such as the peak sensitivity and the threshold. We addressed this using simple mathematical models of RNA thermometers as well as the melt profiles obtained from thermodynamic computations. We found a tradeoff between the thermometer sensitivity and its threshold in a two-state model. For a fixed energy difference between the two states, the threshold was found to be inversely proportional to the peak sensitivity. Moreover, this trend persisted in models with a larger number of states. Furthermore, the fits to the melt profiles, computed based on thermodynamics computations, were consistent with this trend. These results should help to understand the design space of RNA thermometers.

## Results and Discussion

In this study, RNA thermometers were modelled using a two-state model (Fig. 1a., Supp. A.).^14,15^ Such models are used in diverse contexts.^16,17^ The model employed in this study consisted of two states that interconverted into each other at rates *k*_1_ and *k*_2_, respectively. At steady state, the fraction of the unfolded RNA (*y*) was *k*_1_*/*(*k*_1_ + *k*_2_). Since *k*_1_ = *k*_10_ exp (−*E*_1_*/kT*) and *k*_2_ = *k*_20_ exp (−*E*_2_*/kT*), the output fraction was of the form *y*(*T*) = 1*/*(1 + *a* exp (*b/T*)), where *a* = *k*_20_*/k*_10_ and *b* = (*E*_1_ − *E*_2_)*/k*.^18^ In the above expressions, *k* refers to the Boltzmann constant and *T* is the absolute temperature. Furthermore, the parameter sets {*E*_1_, *E*_2_} and {*k*_10_, *k*_20_} model the enthalpic and entropic effects, respectively. Notably, *a* and *b* are both positive, with *b>* 0 since the activation energy of the folded state was assumed to be larger than the activation energy of the unfolded state, *E*_1_ *> E*_2_. In this model, an increase in temperature *T* resulted in an increase in *y*(*T*) as well, as expected from a typical RNA Thermometer response (Fig. 1b.).

**Figure 1.**
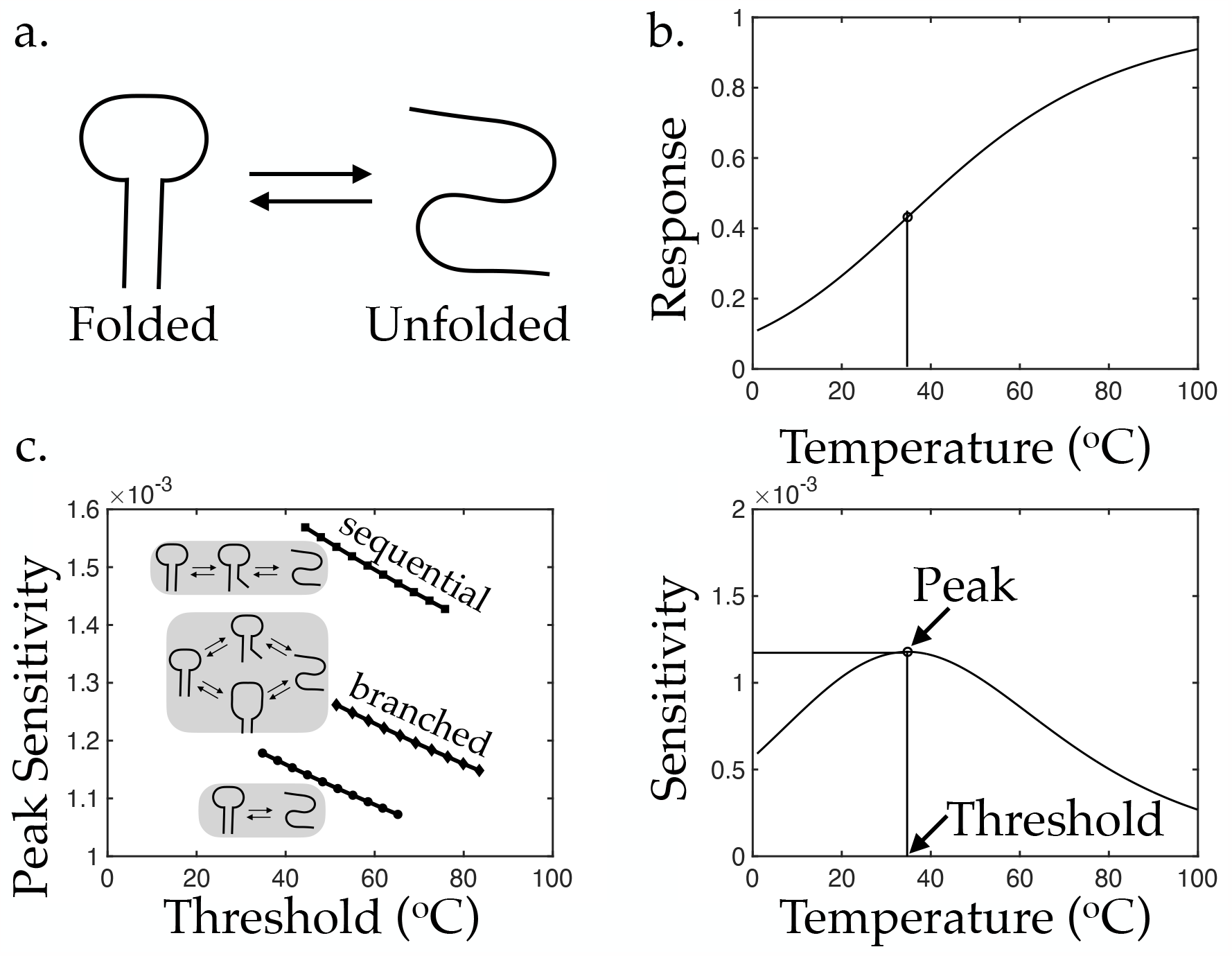
Sensitivity profile in a two-state model and its variants. a. Schematic representation of two states — folded and unfolded — of an RNA thermometer. b. Black lines represent the response of the two-state model (top) and its sensitivity (bottom). The peak sensitivity and threshold temperature are indicated. For simplicity, the temperature axis is shown in °C. Parameters: *a* = 5 × 10^*-*7^, *b* = 4.55 × 10^3^ K. c. Black lines with circles, squares, and diamonds represent the sensitivity profile for the two state model, the sequential variant (*n* +1 = 3), and the branched variant, respectively. Grey boxes show the schematic representations. The parameter *b* was varied in the interval [4.55, 5.00] × 10^3^ K.

The sensitivity of the response was calculated from its derivative (Fig. 1b.), using the following equation:

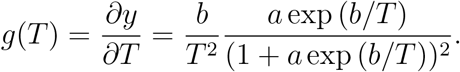

Here, *g*(*T*) was observed to be positive, and it approached 0 as *T* ⟶ 0 or *T* ⟶ ∞. Furthermore, on considering *∂g/∂T*, it was found that *g*(*T*) reached its single maximum value at a temperature between 0 and 1 where *a* exp (*b/T*) = (*b* + 2*T*)*/*(*b −* 2*T*). The maximum value of the sensitivity and the threshold temperature at which this maximum sensitivity was achieved were denoted as *g*_*max*_ and *T*_*threshold*_, respectively. It was found that *T*_*threshold*_ *< b/*2 and that,

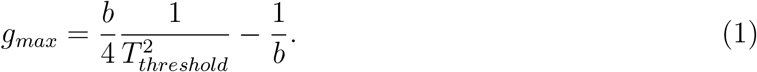

This showed that with an increase in *T*_*threshold*_, for a fixed value of the parameter *b*, the *g*_*max*_ decreased (Fig. 1c.). This indicated a tradeoff between the thermometer sensitivity *g*_*max*_ and its threshold *T*_*threshold*_. Furthermore, on changing the entropic contributions, using parameter *a, g*_*max*_ and *T*_*threshold*_ changed in an inverse proportion. This indicates that modifying *g*_*max*_ for a fixed *T*_*threshold*_, or vice versa, would necessarily need an enthalpic change (the parameter *b*).

From a thermometer design perspective, given the desired *g*_*max*_ and *T*_*threshold*_ values, the parameter *b* could be tuned to meet the *g*_*max*_ specification followed by a tuning of the parameter *a* to meet the *T*_*threshold*_ specification.

To verify the persistence of the above trends in other, more realistic mathematical models that account for multiple folding pathways and states, a sequential model and a branched model were considered.^14,15^ The sequential model comprised a sequence of intermediate states, characterised by progressive unfolding, between the completely folded and the completely unfolded states (Fig. 1c., Supp. B.). A total of *n* +1 states, including the completely folded and the unfolded states, were considered. The branched model contained two reaction paths between the completely folded and the completely unfolded states (Fig. 1c., Supp. C.), symbolized by two separate states with different folding configurations. The equations of this mathematical model were obtained in a similar manner as the two-state model, and computations were performed to generate the respective sensitivity profile *g*(*T*), the maximum sensitivity *g*_*max*_, and the threshold temperature *T*_*threshold*_.

In the sequential *n* + 1-state model, the fraction of the RNA in the completely unfolded state was 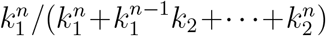. For *k*_1_ = *k*_10_ exp (−*E*_1_*/kT*) and *k*_2_ = *k*_20_ exp (−*E*_2_*/kT*), the output fraction was *y*(*T*) = 1*/*(1 + *a* exp (*b/T*)+ ·· · + *a*^*n*^ exp (*nb/T*)), where *a* = *k*_20_*/k*_10_ and *b* = (*E*_1_ − *E*_2_)*/k*. The sensitivity profile was obtained using the following equation:

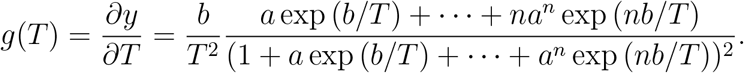

While a general form was not obtained, the *g*_*max*_ and *T*_*threshold*_ were numerically computed from this expression (Fig. 1c.). The plot of *g*_*max*_ versus *T*_*threshold*_ was overlaid with the previously obtained plot from the two-state model. These plots were similar for the parameter sets considered.

In the case of the branched model, the fraction of the RNA in the completely unfolded state was found to be (*k*_1_*/*(*k*_1_ + *k*_2_))^2^. For *k*_1_ and *k*_2_, as defined above, the output fraction was *y*(*T*) = 1*/*(1 + *a* exp (*b/T*))^2^. Therefore, the sensitivity profile could be expressed as

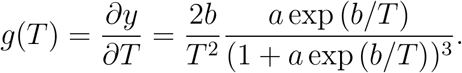

Similar to the sequential model, the *g*_*max*_ and *T*_*threshold*_ were computed from this expression. It was noted that, for the parameters considered, the plots of *y*_*max*_ versus *T*_1*/*2_ and *g*_*max*_ versus *T*_*threshold*_ exhibited a similar trend as the sequential model (Fig. 1c.).

Furthermore, to verify the persistence of the above constraints in even more realistic models, we used NUPACK — a webserver for thermodynamic computations of nucleic acids.^19^ The melt profiles of a library of RNA thermometers, obtained from variations of a previously constructed synthetic RNA thermometer,^8,10^ were computed and analyzed. The set of RNA thermometers were obtained by mutating a single base of the starting RNA thermometer (Supp. D.). A melt profile yields the probability of a base being unpaired at a specific temperature. For each variant thermometer, we computed the melt profile of the RBS, by averaging the melt profiles of each base in the RBS (Fig. 2a, Supp. E.). These melt profiles were computed from 1 °C to 100 °C at a resolution of 1 °C.

**Figure 2.**
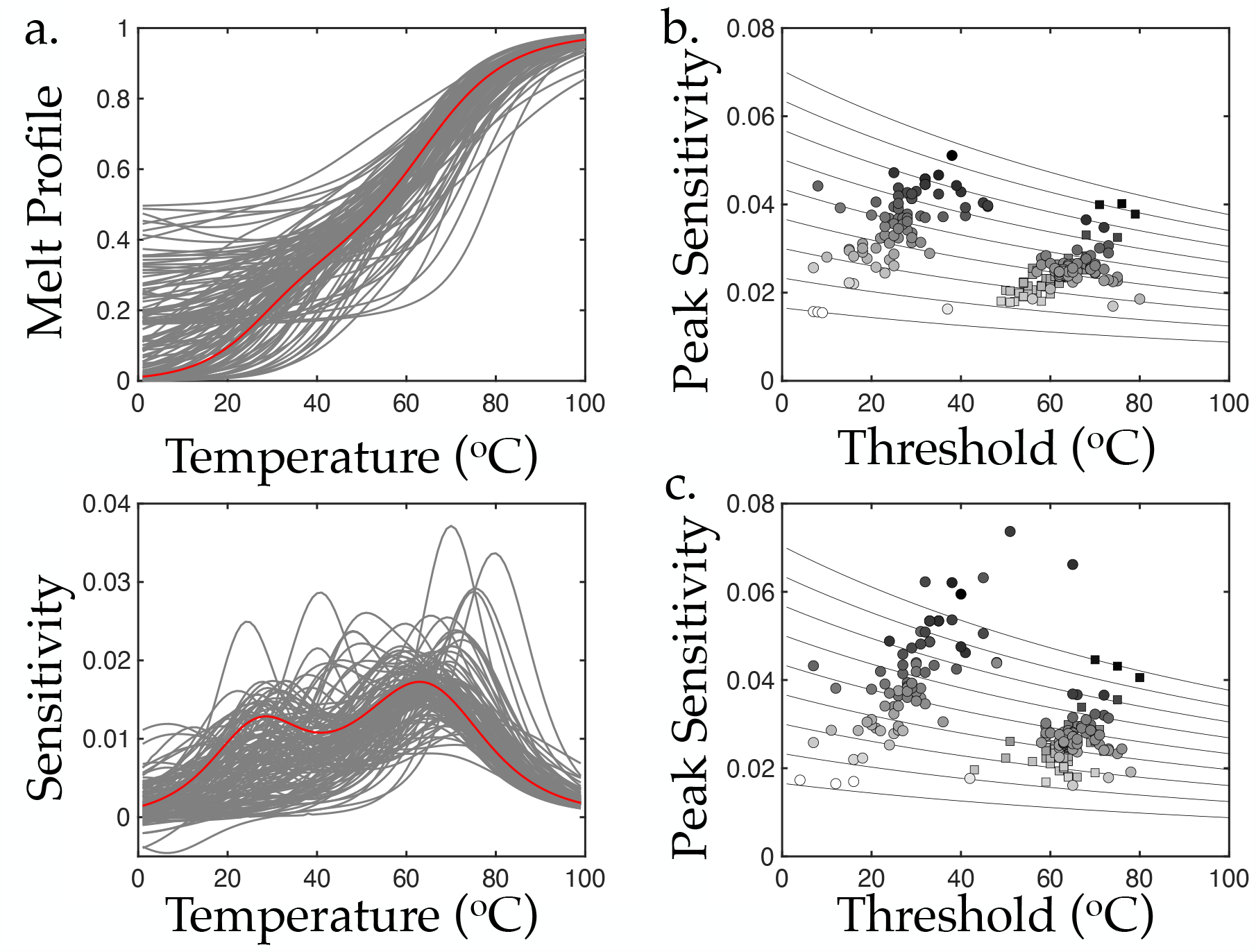
Tradeoffs in computed melt profiles. a. Red line represents the computed melt profile and sensitivity of an RNA thermometer. Grey lines represent the computed melt profiles and sensitivities of a library of RNA thermometers generated from the starting thermometer. b. Circles and squares represent the peak sensitivity and threshold of fits to the melt profiles having one and two peaks, respectively. The fill colour represents the value of the parameter *b*. Thin black lines represent a grid of lines generated from Eqn. (1) for different values of *b*. c. Same as in b., with the peak sensitivity and threshold coming from the NUPACK-generated melt profile rather than the fit.

Subsequently, the peak sensitivity and the threshold temperature of the response were computed using the derivative of the melt profiles (Supp. F.). A plot of the peak sensitivity versus the threshold showed two clusters (Fig. 2c), resulting from the presence of two or more peaks in the melt profiles, possibly due to the melting of multiple substructures. However, a direct examination of the possible sensitivity-threshold tradeoff was challenging because multiple underlying parameters could have changed when the identity of the base was modified. Therefore, the melt profiles were fitted to various functional forms to identify the presence of tradeoffs.

First, the melt profiles were fitted to a parametric function, as in the two-state model (Supp. G.). Since the derivative of some melt profiles exhibited two peaks, these specific melt profiles were fitted to a sum of two functions. These fits were found to work better, principally because this function allowed for a better fit in terms of the the two peaks. The peak sensitivities of the melt profile fits were plotted against their threshold values, and then segregated according to the *b* value (Fig. 2b). They accurately satisfied the sensitivity-threshold tradeoffs, as anticipated from the above analysis of the two-state model and its variants. Furthermore, even the senstivity-threshold plots obtained from the NUPACK melt profiles, similarly segregated, were largely consistent with the sensitivity-threshold tradeoffs (Fig. 2c). The relatively minor deviations could have resulted from errors in the fits, owing to higher order dynamics in the underlying thermodynamic model. We concluded that the parameters obtained from the fits to the melt profiles were consistent with the sensitivity-threshold tradeoffs.

In view of the fact that RNA thermometers can mediate cellular responses to temperature, we inverstigated whether there are constraints that limit the achievable sensitivities and thresholds in the temperature response of RNA thermometers. Using a two-state model, we identified a tradeoff in the sensitivity and threshold of RNA thermometers. Moreover, this trend persisted in larger models of RNA thermometers as well. We computed the melt profiles of synthetic RNA thermometers, using the thermodynamics-based webserver NUPACK, finding that these were consistent with the above constraints.

## Supporting Information Available

The following files are available free of charge.

- Supplementary: A. Two-State Model, B. Sequential Model, C. Branched Model, D. RNA Thermometer Sequences, E. NUPACK Melt Profiles, F. Determination of Peak Sensitivity and Threshold, G. Melt Profile Fits.

## Notes

### Competing Interest Statement

The authors have declared no competing interest.

