## Supplementary Information for "Tradeoffs in the Design of RNA Thermometers"

### Contents

A. Two-State Model (Pages 3)

B. Sequential Model (Page 4)

C. Branched Model (Pages 5)

D. RNA Thermometer Sequences (Pages 6-12)

E. NUPACK Melt Profiles (Page 13)

F. Determination of Peak Sensitivity and Threshold (Page 14)

G. Melt Profile Fits (Pages 15-23)

#### A. Two-State Model

- Steady state

At steady state, $k_1[\text{folded}] = k_2[\text{unfolded}]$.

$$\Rightarrow y = \frac{[\text{folded}]}{[\text{folded}] + [\text{unfolded}]} = \frac{k_1}{k_1 + k_2}.$$

Using $k_1 = k_{10}\exp(-E_1/kT)$, $k_2 = k_{20}\exp(-E_2/kT)$,

$$y = \frac{1}{1 + a\exp(b/T)},$$

where $a = k_{20}/k_{10}$ and $b = (E_1 - E_2)/k$.

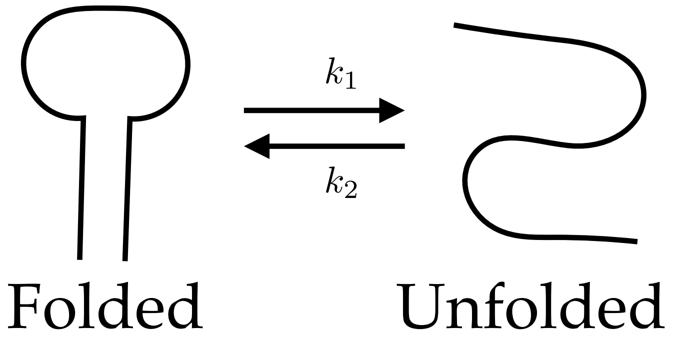

- Sensitivity

$$g(T) = \frac{\partial y}{\partial T} = \frac{b}{T^2}\frac{a\exp(b/T)}{(1 + a\exp(b/T))^2}$$

Properties:

1. $g(T) > 0$ (from expression),
2. $\lim_{T\to 0} g(T) = 0$ (L'Hôpital's rule),
3. $\lim_{T\to\infty} g(T) = 0$ (L'Hôpital's rule).

- Sensitivity is maximum when

$$\frac{\partial g}{\partial T} = 0 \Rightarrow a\exp(b/T) = \frac{b + 2T}{b - 2T}.$$

Denote maximizing $T$ as $T_{threshold}$,
and maxima as $g_{max} = g(T_{threshold})$.

$$\Rightarrow g_{max} = \frac{b}{T_{threshold}^2}\frac{\frac{b+2T_{threshold}}{b-2T_{threshold}}}{(1 + \frac{b+2T_{threshold}}{b-2T_{threshold}})^2},$$

$$\Rightarrow g_{max} = \frac{b}{4}\frac{1}{T_{threshold}^2} - \frac{1}{b}.$$

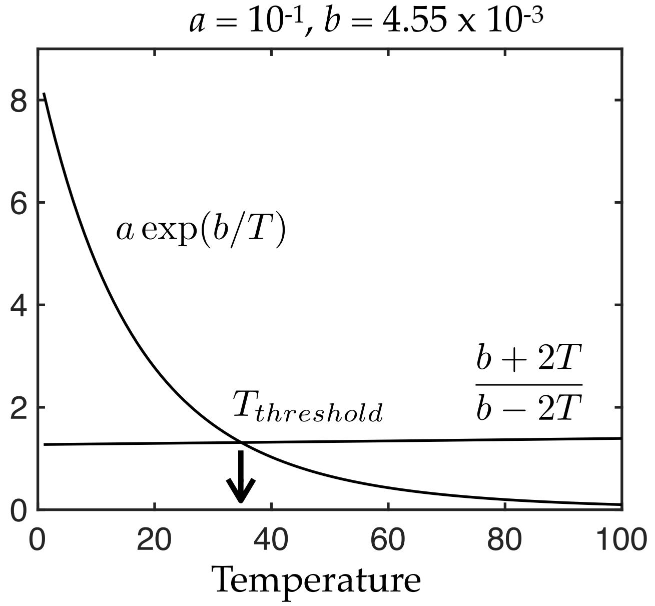

#### B. Sequential Model

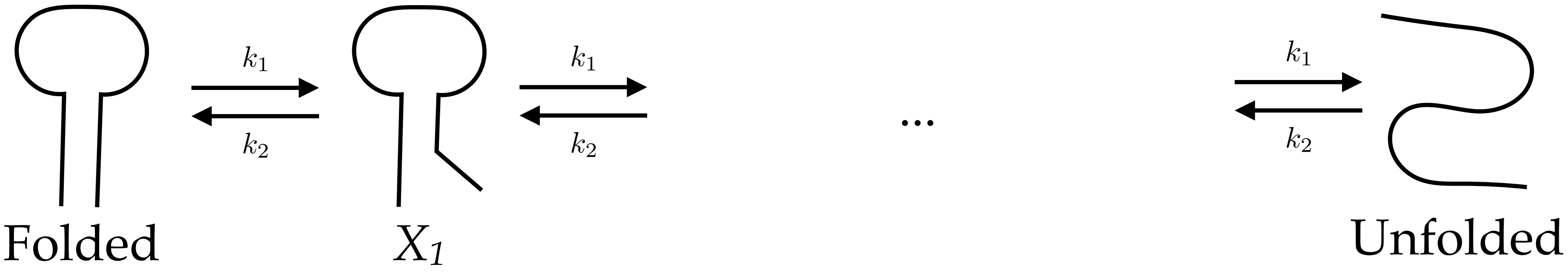

- Steady state

At steady state, $k_1[\text{folded}] = k_2[X_1], \cdots k_1[X_{n-1}] = k_2[\text{unfolded}]$.

$$\Rightarrow y = \frac{[\text{folded}]}{[\text{folded}] + [X_1] + \ldots + [\text{unfolded}]} = \frac{k_1^n}{k_1^n + k_1^{n-1}k_2 + \ldots k_2^n}.$$

Using $k_1 = k_{10}\exp(-E_1/kT)$, $k_2 = k_{20}\exp(-E_2/kT)$,

$$y = \frac{1}{1 + a\exp\left(b/T\right) + \ldots \left(a\exp\left(b/T\right)\right)^n},$$

where $a = k_{20}/k_{10}$ and $b = (E_1 - E_2)/k$.

- Sensitivity

$$g(T) = \frac{\partial y}{\partial T} = \frac{b}{T^2}\frac{a\exp\left(b/T\right) + \cdots + na^n\exp\left(nb/T\right)}{\left(1 + a\exp\left(b/T\right) + \cdots + a^n\exp\left(nb/T\right)\right)^2}.$$

- $n+1 = 3$ is considered in the main text.

#### C. Branched Model

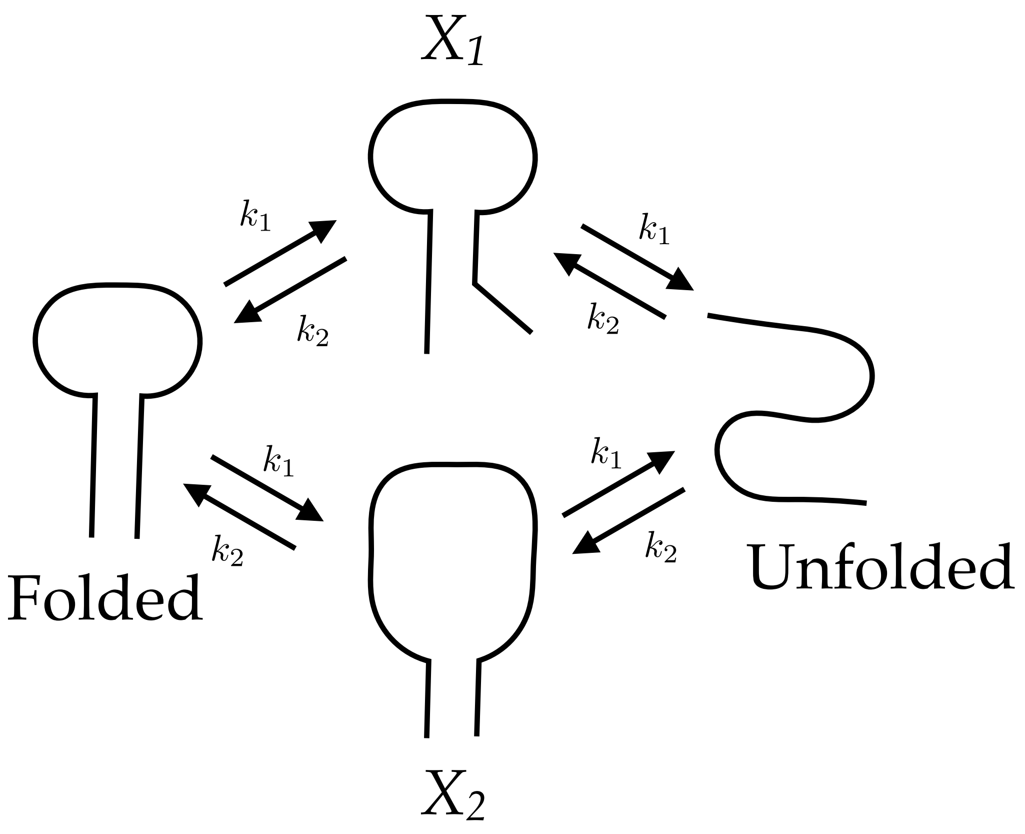

- Steady state

At steady state,
$k_1[\text{folded}] = k_2[X_1]$, $k_1[X_1] = k_2[\text{unfolded}]$,
$k_1[\text{folded}] = k_2[X_2]$, $k_1[X_2] = k_2[\text{unfolded}]$.

$$\Rightarrow y = \frac{[\text{folded}]}{[\text{folded}] + [X_1] + [X_2] + [\text{unfolded}]} = \left(\frac{k_1}{k_1 + k_2}\right)^2.$$

Using $k_1 = k_{10}\exp(-E_1/kT)$, $k_2 = k_{20}\exp(-E_2/kT)$,

$$y = \left(\frac{1}{1 + a\exp(b/T)}\right)^2.$$

where $a = k_{20}/k_{10}$ and $b = (E_1 - E_2)/k$.

- Sensitivity

$$g(T) = \frac{\partial y}{\partial T} = \frac{2b}{T^2}\frac{a\exp(b/T)}{(1 + a\exp(b/T))^3}.$$

#### D. RNA Thermometer Sequences

Red text is the RBS. Blue text denotes the base change or deletion.

| # | Name | Sequence |
|---|---|---|
| 0 | X | GGAUCCCUCACUUACUAGUCUGCAGAAGGAGAUAUACCCAUGG |
| 1 | 1O | -GAUCCCUCACUUACUAGUCUGCAGAAGGAGAUAUACCCAUGG |
| 2 | 1A | AGAUCCCUCACUUACUAGUCUGCAGAAGGAGAUAUACCCAUGG |
| 3 | 1C | CGAUCCCUCACUUACUAGUCUGCAGAAGGAGAUAUACCCAUGG |
| 4 | 1U | UGAUCCCUCACUUACUAGUCUGCAGAAGGAGAUAUACCCAUGG |
| 5 | 2O | G-AUCCCUCACUUACUAGUCUGCAGAAGGAGAUAUACCCAUGG |
| 6 | 2A | GAAUCCCUCACUUACUAGUCUGCAGAAGGAGAUAUACCCAUGG |
| 7 | 2C | GCAUCCCUCACUUACUAGUCUGCAGAAGGAGAUAUACCCAUGG |
| 8 | 2U | GUAUCCCUCACUUACUAGUCUGCAGAAGGAGAUAUACCCAUGG |
| 9 | 3O | GG-UCCCUCACUUACUAGUCUGCAGAAGGAGAUAUACCCAUGG |
| 10 | 3C | GGCUCCCUCACUUACUAGUCUGCAGAAGGAGAUAUACCCAUGG |
| 11 | 3G | GGGUCCCUCACUUACUAGUCUGCAGAAGGAGAUAUACCCAUGG |
| 12 | 3U | GGUUCCCUCACUUACUAGUCUGCAGAAGGAGAUAUACCCAUGG |
| 13 | 4O | GGA-CCCUCACUUACUAGUCUGCAGAAGGAGAUAUACCCAUGG |
| 14 | 4A | GGAACCCUCACUUACUAGUCUGCAGAAGGAGAUAUACCCAUGG |
| 15 | 4C | GGACCCCUCACUUACUAGUCUGCAGAAGGAGAUAUACCCAUGG |
| 16 | 4G | GGAGCCCUCACUUACUAGUCUGCAGAAGGAGAUAUACCCAUGG |
| 17 | 5O | GGAU-CCUCACUUACUAGUCUGCAGAAGGAGAUAUACCCAUGG |
| 18 | 5A | GGAUACCUCACUUACUAGUCUGCAGAAGGAGAUAUACCCAUGG |
| 19 | 5G | GGAUGCCUCACUUACUAGUCUGCAGAAGGAGAUAUACCCAUGG |
| 20 | 5U | GGAUUCCUCACUUACUAGUCUGCAGAAGGAGAUAUACCCAUGG |

Red text is the RBS. Blue text denotes the base change or deletion.

| # | Name | Sequence |
|---|---|---|
| 0 | X | GGAUCCCUCACUUACUAGUCUGCAGAAGGAGAUAUACCCAUGG |
| 21 | 6O | GGAUC-CUCACUUACUAGUCUGCAGAAGGAGAUAUACCCAUGG |
| 22 | 6A | GGAUCACUCACUUACUAGUCUGCAGAAGGAGAUAUACCCAUGG |
| 23 | 6G | GGAUCGCUCACUUACUAGUCUGCAGAAGGAGAUAUACCCAUGG |
| 24 | 6U | GGAUCUCUCACUUACUAGUCUGCAGAAGGAGAUAUACCCAUGG |
| 25 | 7O | GGAUCC-UCACUUACUAGUCUGCAGAAGGAGAUAUACCCAUGG |
| 26 | 7A | GGAUCCAUCACUUACUAGUCUGCAGAAGGAGAUAUACCCAUGG |
| 27 | 7G | GGAUCCGUCACUUACUAGUCUGCAGAAGGAGAUAUACCCAUGG |
| 28 | 7U | GGAUCCUUCACUUACUAGUCUGCAGAAGGAGAUAUACCCAUGG |
| 29 | 8O | GGAUCCC-CACUUACUAGUCUGCAGAAGGAGAUAUACCCAUGG |
| 30 | 8A | GGAUCCCACACUUACUAGUCUGCAGAAGGAGAUAUACCCAUGG |
| 31 | 8C | GGAUCCCCCACUUACUAGUCUGCAGAAGGAGAUAUACCCAUGG |
| 32 | 8G | GGAUCCCGCACUUACUAGUCUGCAGAAGGAGAUAUACCCAUGG |
| 33 | 9O | GGAUCCCU-ACUUACUAGUCUGCAGAAGGAGAUAUACCCAUGG |
| 34 | 9A | GGAUCCCUAACUUACUAGUCUGCAGAAGGAGAUAUACCCAUGG |
| 35 | 9G | GGAUCCCUGACUUACUAGUCUGCAGAAGGAGAUAUACCCAUGG |
| 36 | 9U | GGAUCCCUUACUUACUAGUCUGCAGAAGGAGAUAUACCCAUGG |
| 37 | 10O | GGAUCCCUC-CUUACUAGUCUGCAGAAGGAGAUAUACCCAUGG |
| 38 | 10C | GGAUCCCUCCCUUACUAGUCUGCAGAAGGAGAUAUACCCAUGG |
| 39 | 10G | GGAUCCCUCGCUUACUAGUCUGCAGAAGGAGAUAUACCCAUGG |
| 40 | 10U | GGAUCCCUCUCUUACUAGUCUGCAGAAGGAGAUAUACCCAUGG |

Red text is the RBS. Blue text denotes the base change or deletion.

| # | Name | Sequence |
|---|---|---|
| 0 | X | GGAUCCCUCACUUACUAGUCUGCAGAAGGAGAUAUACCCAUGG |
| 41 | 11O | GGAUCCCUCA-UUACUAGUCUGCAGAAGGAGAUAUACCCAUGG |
| 42 | 11A | GGAUCCCUCAAUUACUAGUCUGCAGAAGGAGAUAUACCCAUGG |
| 43 | 11G | GGAUCCCUCAGUUACUAGUCUGCAGAAGGAGAUAUACCCAUGG |
| 44 | 11U | GGAUCCCUCAUUUACUAGUCUGCAGAAGGAGAUAUACCCAUGG |
| 45 | 12O | GGAUCCCUCAC-UACUAGUCUGCAGAAGGAGAUAUACCCAUGG |
| 46 | 12A | GGAUCCCUCACAUACUAGUCUGCAGAAGGAGAUAUACCCAUGG |
| 47 | 12C | GGAUCCCUCACCUACUAGUCUGCAGAAGGAGAUAUACCCAUGG |
| 48 | 12G | GGAUCCCUCACGUACUAGUCUGCAGAAGGAGAUAUACCCAUGG |
| 49 | 13O | GGAUCCCUCACU-ACUAGUCUGCAGAAGGAGAUAUACCCAUGG |
| 50 | 13A | GGAUCCCUCACUAACUAGUCUGCAGAAGGAGAUAUACCCAUGG |
| 51 | 13C | GGAUCCCUCACUCACUAGUCUGCAGAAGGAGAUAUACCCAUGG |
| 52 | 13G | GGAUCCCUCACUGACUAGUCUGCAGAAGGAGAUAUACCCAUGG |
| 53 | 14O | GGAUCCCUCACUU-CUAGUCUGCAGAAGGAGAUAUACCCAUGG |
| 54 | 14C | GGAUCCCUCACUUCCUAGUCUGCAGAAGGAGAUAUACCCAUGG |
| 55 | 14G | GGAUCCCUCACUUGCUAGUCUGCAGAAGGAGAUAUACCCAUGG |
| 56 | 14U | GGAUCCCUCACUUUCUAGUCUGCAGAAGGAGAUAUACCCAUGG |
| 57 | 15O | GGAUCCCUCACUUA-UAGUCUGCAGAAGGAGAUAUACCCAUGG |
| 58 | 15A | GGAUCCCUCACUUAAUAGUCUGCAGAAGGAGAUAUACCCAUGG |
| 59 | 15G | GGAUCCCUCACUUAGUAGUCUGCAGAAGGAGAUAUACCCAUGG |
| 60 | 15U | GGAUCCCUCACUUAUUAGUCUGCAGAAGGAGAUAUACCCAUGG |

Red text is the RBS. Blue text denotes the base change or deletion.

| # | Name | Sequence |
|---|---|---|
| 0 | X | GGAUCCCUCACUUACUAGUCUGCAGAAGGAGAUAUACCCAUGG |
| 61 | 160 | GGAUCCCUCACUUAC-AGUCUGCAGAAGGAGAUAUACCCAUGG |
| 62 | 16A | GGAUCCCUCACUUACAAGUCUGCAGAAGGAGAUAUACCCAUGG |
| 63 | 16C | GGAUCCCUCACUUACCAGUCUGCAGAAGGAGAUAUACCCAUGG |
| 64 | 16G | GGAUCCCUCACUUACGAGUCUGCAGAAGGAGAUAUACCCAUGG |
| 65 | 170 | GGAUCCCUCACUUACU-GUCUGCAGAAGGAGAUAUACCCAUGG |
| 66 | 17C | GGAUCCCUCACUUACUCGUCUGCAGAAGGAGAUAUACCCAUGG |
| 67 | 17G | GGAUCCCUCACUUACUGGUCUGCAGAAGGAGAUAUACCCAUGG |
| 68 | 17U | GGAUCCCUCACUUACUUGUCUGCAGAAGGAGAUAUACCCAUGG |
| 69 | 180 | GGAUCCCUCACUUACUA-UCUGCAGAAGGAGAUAUACCCAUGG |
| 70 | 18A | GGAUCCCUCACUUACUAAUCUGCAGAAGGAGAUAUACCCAUGG |
| 71 | 18C | GGAUCCCUCACUUACUACUCUGCAGAAGGAGAUAUACCCAUGG |
| 72 | 18U | GGAUCCCUCACUUACUAUUCUGCAGAAGGAGAUAUACCCAUGG |
| 73 | 190 | GGAUCCCUCACUUACUAG-CUGCAGAAGGAGAUAUACCCAUGG |
| 74 | 19A | GGAUCCCUCACUUACUAGACUGCAGAAGGAGAUAUACCCAUGG |
| 75 | 19C | GGAUCCCUCACUUACUAGCCUGCAGAAGGAGAUAUACCCAUGG |
| 76 | 19G | GGAUCCCUCACUUACUAGGCUGCAGAAGGAGAUAUACCCAUGG |
| 77 | 200 | GGAUCCCUCACUUACUAGU-UGCAGAAGGAGAUAUACCCAUGG |
| 78 | 20A | GGAUCCCUCACUUACUAGUAUGCAGAAGGAGAUAUACCCAUGG |
| 79 | 20G | GGAUCCCUCACUUACUAGUGUGCAGAAGGAGAUAUACCCAUGG |
| 80 | 20U | GGAUCCCUCACUUACUAGUUUGCAGAAGGAGAUAUACCCAUGG |

Red text is the RBS. Blue text denotes the base change or deletion.

| # | Name | Sequence |
|---|---|---|
| **0** | X | GGAUCCCUCACUUACUAGUCUGCAGAAGGAGAUAUACCCAUGG |
| 81 | 210 | GGAUCCCUCACUUACUAGUC-GCAGAAGGAGAUAUACCCAUGG |
| 82 | 21A | GGAUCCCUCACUUACUAGUCAGCAGAAGGAGAUAUACCCAUGG |
| 83 | 21C | GGAUCCCUCACUUACUAGUCCGCAGAAGGAGAUAUACCCAUGG |
| 84 | 21G | GGAUCCCUCACUUACUAGUCGGCAGAAGGAGAUAUACCCAUGG |
| 85 | 220 | GGAUCCCUCACUUACUAGUCU-CAGAAGGAGAUAUACCCAUGG |
| 86 | 22A | GGAUCCCUCACUUACUAGUCUACAGAAGGAGAUAUACCCAUGG |
| 87 | 22C | GGAUCCCUCACUUACUAGUCUCCAGAAGGAGAUAUACCCAUGG |
| 88 | 22U | GGAUCCCUCACUUACUAGUCUUCAGAAGGAGAUAUACCCAUGG |
| 89 | 230 | GGAUCCCUCACUUACUAGUCUG-AGAAGGAGAUAUACCCAUGG |
| 90 | 23A | GGAUCCCUCACUUACUAGUCUGAAGAAGGAGAUAUACCCAUGG |
| 91 | 23G | GGAUCCCUCACUUACUAGUCUGGAGAAGGAGAUAUACCCAUGG |
| 92 | 23U | GGAUCCCUCACUUACUAGUCUGUAGAAGGAGAUAUACCCAUGG |
| 93 | 240 | GGAUCCCUCACUUACUAGUCUGC-GAAGGAGAUAUACCCAUGG |
| 94 | 24C | GGAUCCCUCACUUACUAGUCUGCCGAAGGAGAUAUACCCAUGG |
| 95 | 24G | GGAUCCCUCACUUACUAGUCUGCGGAAGGAGAUAUACCCAUGG |
| 96 | 24U | GGAUCCCUCACUUACUAGUCUGCUGAAGGAGAUAUACCCAUGG |
| 97 | 250 | GGAUCCCUCACUUACUAGUCUGCA-AAGGAGAUAUACCCAUGG |
| 98 | 25A | GGAUCCCUCACUUACUAGUCUGCAAAAGGAGAUAUACCCAUGG |
| 99 | 25C | GGAUCCCUCACUUACUAGUCUGCACAAGGAGAUAUACCCAUGG |
| 100 | 25U | GGAUCCCUCACUUACUAGUCUGCAUAAGGAGAUAUACCCAUGG |

Red text is the RBS. Blue text denotes the base change or deletion.

| # | Name | Sequence |
|---|---|---|
| **0** | **X** | GGAUCCCUCACUUACUAGUCUGCAGAAGGAGAUAUACCCAUGG |
| 101 | 320 | GGAUCCCUCACUUACUAGUCUGCAGAAGGAG-UAUACCCAUGG |
| 102 | 32C | GGAUCCCUCACUUACUAGUCUGCAGAAGGAGCUAUACCCAUGG |
| 103 | 32G | GGAUCCCUCACUUACUAGUCUGCAGAAGGAGGUAUACCCAUGG |
| 104 | 32U | GGAUCCCUCACUUACUAGUCUGCAGAAGGAGUUAUACCCAUGG |
| 105 | 330 | GGAUCCCUCACUUACUAGUCUGCAGAAGGAGA-AUACCCAUGG |
| 106 | 33A | GGAUCCCUCACUUACUAGUCUGCAGAAGGAGAAAUACCCAUGG |
| 107 | 33C | GGAUCCCUCACUUACUAGUCUGCAGAAGGAGACAUACCCAUGG |
| 108 | 33G | GGAUCCCUCACUUACUAGUCUGCAGAAGGAGAGAUACCCAUGG |
| 109 | 340 | GGAUCCCUCACUUACUAGUCUGCAGAAGGAGAU-UACCCAUGG |
| 110 | 34C | GGAUCCCUCACUUACUAGUCUGCAGAAGGAGAUCUACCCAUGG |
| 111 | 34G | GGAUCCCUCACUUACUAGUCUGCAGAAGGAGAUGUACCCAUGG |
| 112 | 34U | GGAUCCCUCACUUACUAGUCUGCAGAAGGAGAUUUACCCAUGG |
| 113 | 350 | GGAUCCCUCACUUACUAGUCUGCAGAAGGAGAUU-ACCCAUGG |
| 114 | 35A | GGAUCCCUCACUUACUAGUCUGCAGAAGGAGAUAAACCCAUGG |
| 115 | 35C | GGAUCCCUCACUUACUAGUCUGCAGAAGGAGAUACACCCAUGG |
| 116 | 35G | GGAUCCCUCACUUACUAGUCUGCAGAAGGAGAUAGACCCAUGG |
| 117 | 360 | GGAUCCCUCACUUACUAGUCUGCAGAAGGAGAUAU-CCCAUGG |
| 118 | 36C | GGAUCCCUCACUUACUAGUCUGCAGAAGGAGAUAUCCCCAUGG |
| 119 | 36G | GGAUCCCUCACUUACUAGUCUGCAGAAGGAGAUAUGCCCAUGG |
| 120 | 36U | GGAUCCCUCACUUACUAGUCUGCAGAAGGAGAUAUUCCCAUGG |

Red text is the RBS. Blue text denotes the base change or deletion.

| # | Name | Sequence |
|---|---|---|
| **0** | **X** | GGAUCCCUCACUUACUAGUCUGCAGAAGGAGAUAUACCCAUGG |
| 121 | 370 | GGAUCCCUCACUUACUAGUCUGCAGAAGGAGAUAUA-CCAUGG |
| 122 | 37A | GGAUCCCUCACUUACUAGUCUGCAGAAGGAGAUAUAACCAUGG |
| 123 | 37G | GGAUCCCUCACUUACUAGUCUGCAGAAGGAGAUAUAGCCAUGG |
| 124 | 37U | GGAUCCCUCACUUACUAGUCUGCAGAAGGAGAUAUAUCCAUGG |
| 125 | 380 | GGAUCCCUCACUUACUAGUCUGCAGAAGGAGAUAUAC-CAUGG |
| 126 | 38A | GGAUCCCUCACUUACUAGUCUGCAGAAGGAGAUAUACACAUGG |
| 127 | 38G | GGAUCCCUCACUUACUAGUCUGCAGAAGGAGAUAUACGCAUGG |
| 128 | 38U | GGAUCCCUCACUUACUAGUCUGCAGAAGGAGAUAUACUCAUGG |
| 129 | 390 | GGAUCCCUCACUUACUAGUCUGCAGAAGGAGAUAUACC-AUGG |
| 130 | 39A | GGAUCCCUCACUUACUAGUCUGCAGAAGGAGAUAUACCAAUGG |
| 131 | 39G | GGAUCCCUCACUUACUAGUCUGCAGAAGGAGAUAUACCGAUGG |
| 132 | 39U | GGAUCCCUCACUUACUAGUCUGCAGAAGGAGAUAUACCUAUGG |
| 133 | 400 | GGAUCCCUCACUUACUAGUCUGCAGAAGGAGAUAUACCC-UGG |
| 134 | 40C | GGAUCCCUCACUUACUAGUCUGCAGAAGGAGAUAUACCCCUGG |
| 135 | 40G | GGAUCCCUCACUUACUAGUCUGCAGAAGGAGAUAUACCCGUGG |
| 136 | 40U | GGAUCCCUCACUUACUAGUCUGCAGAAGGAGAUAUACCCUUGG |

#### E. NUPACK Melt Profiles

a. Melt profiles obtained from NUPACK. Red line is the melt profile of the starting thermometer. Grey lines are the melt profiles of the other thermometers. b. Melt profiles obtained by averaging the probabilities that an element of the RBS was unpaired. This is same as in Fig. 2a. of main text.

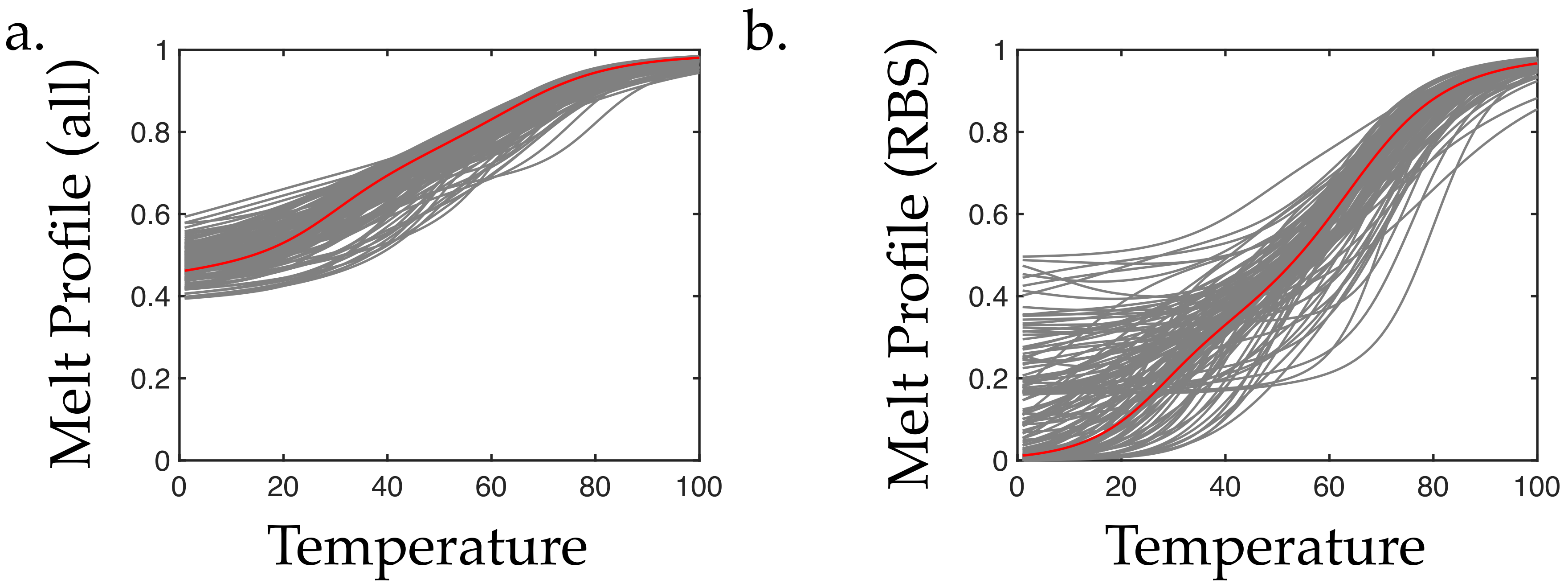

#### F. Determination of Peak Sensitivity and Threshold

The derivative of the melt profile was obtained using the first difference. the peak sensitivities and thresholds were obtained using the MATLAB function 'findpeaks'. Peaks with width > 10 $^{o}$C were considered for analysis. Out of the 136 sensitivity profiles, 56, 78, and 2 had one peak, two peaks, and three peaks, respectively. Blue squares, red circles, and black diamonds represent the thermometers with one peak, two peaks, and three peaks, respectively.

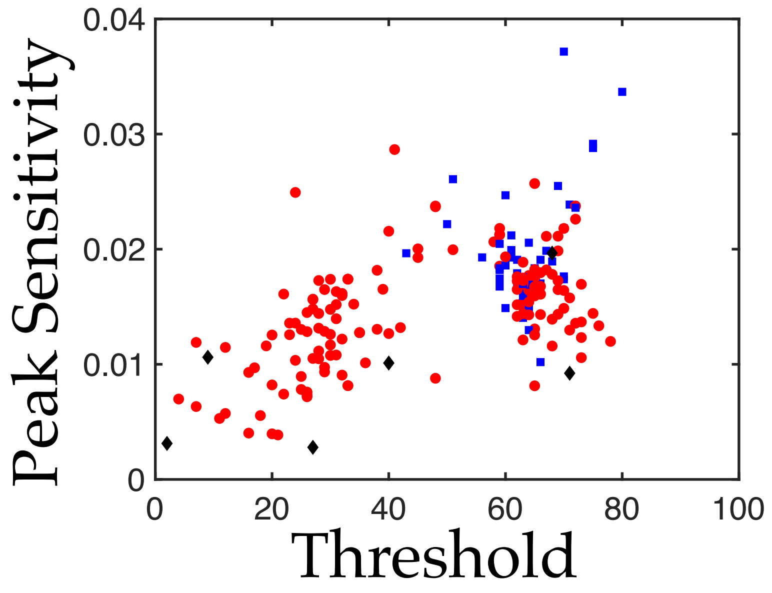

#### G. Melt Profile Fits

- The model used to fit to a melt profile with one peak is

$$y(T) = \frac{x_3}{1+\exp(x_2(\frac{1}{T+273}-\frac{1}{x_1}))} + (1-x_3),$$

where $T$ is the input temperature (in $^{o}$C), y is the response output, and [$x_1$, $x_2$, $x_3$] are parameters.
- The method use for fitting is least-squares, implemented as the MATLAB function 'lsqcurvefit' with initial conditions = [323, 5000, $10^{-1}$], lower bound = [273, 0, $10^{-2}$], and upper bound = [373, 20000, 1].

- The model used to fit to a melt profile with two peaks is

$$y(T) = \frac{x_3}{1+\exp(x_2(\frac{1}{T+273}-\frac{1}{x_1}))} + \frac{x_6}{1+\exp(x_5(\frac{1}{T+273}-\frac{1}{x_4}))} + (1-x_3-x_6),$$

where $T$ is the input temperature (in $^{o}$C), y is the response output, and [$x_1$, $x_2$, $x_3$, $x_4$, $x_5$, $x_6$] are parameters.
- The method use for fitting is least-squares, implemented as the MATLAB function 'lsqcurvefit' with with initial conditions = [285, 4000, $10^{-5}$, 330, 6000, $10^{-1}$], lower bound = [273, 5000, $10^{-2}$, 273, 5000, $10^{-2}$], and upper bound = [373, 20000, 1, 373, 20000, 1].

- Sample fits for melt profiles with one peak and two peak are show in Page 16.
- The fitted parameters for melt profiles with one peak are shown in Pages 17-19.
- The fitted parameters for melt profiles with two peaks are shown in Pages 20-23.

- The peak sensitivities in Fig. 2b. and Fig. 2c. corresponding to a melt profile with one peak were normalized by the parameter $x_3$.
- The peak sensitivities in Fig. 2b. and Fig. 2c. corresponding to a melt profiles with two peaks were normalized by the parameter $x_3$ or $x_6$, depending on whether they corresponded to the first term or the second term, respectively.

### Sample fits

a. Black line is the NUPACK melt profile of Thermometer #135. Red line is the fit. b. Black line is the sensitivity profile of Thermometer #135. Red line is the sensitivity profile of its fit. Black and red circles indicate the peak sensitivity and threshold of the melt profiles and its fit, respectively. c. Black line is the NUPACK melt profile of Thermometer #0. Red line is the fit. b. Black line is the sensitivity profile of Thermometer #0. Red line is the sensitivity profile of its fit. Black and red circles indicate the peak sensitivity and threshold of the melt profiles and its fit, respectively. Green and blue lines represent the derivatives of the individual functions in the fit.

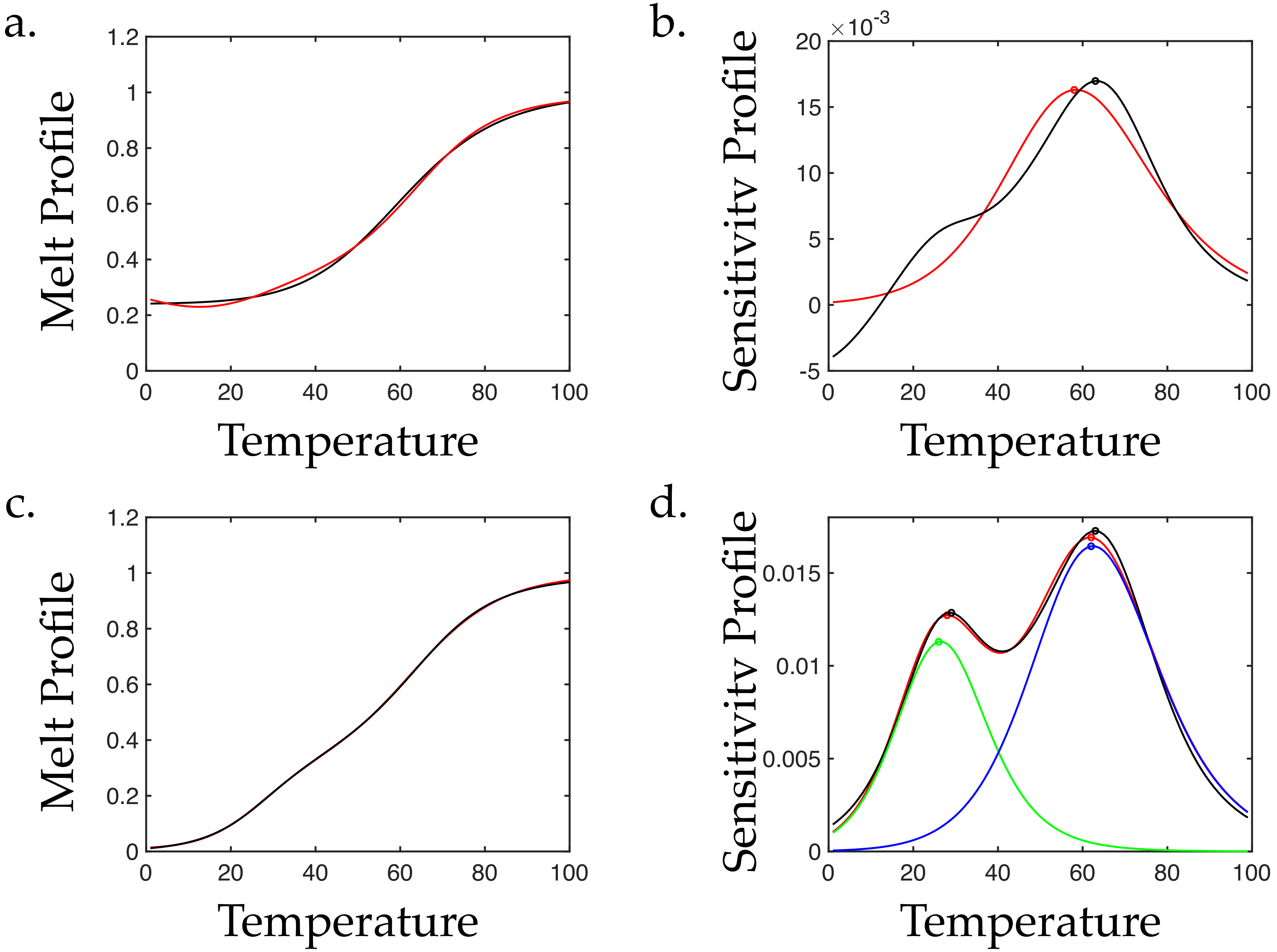

$$y(T) = \frac{x_3}{1 + \exp(x_2(\frac{1}{T+273} - \frac{1}{x_1}))} + (1 - x_3),$$

| # | Thermometer # | $x_1$ | $x_2$ | $x_3$ |
|---|---|---|---|---|
| 1 | 1 | 332.3841281 | 8730.506023 | 0.776624875 |
| 2 | 2 | 333.5938121 | 9604.320563 | 0.774017723 |
| 3 | 3 | 333.1563951 | 9534.299358 | 0.789881457 |
| 4 | 4 | 334.412368 | 9838.596854 | 0.748700471 |
| 5 | 5 | 332.3841281 | 8730.506023 | 0.776624875 |
| 6 | 6 | 334.7049159 | 10081.02001 | 0.747482746 |
| 7 | 7 | 334.8160424 | 10025.76205 | 0.722088542 |
| 8 | 8 | 328.6977103 | 8571.923753 | 0.983679302 |
| 9 | 11 | 331.0994271 | 10535.35433 | 0.999997818 |
| 10 | 13 | 328.3531145 | 7692.847477 | 0.823820823 |
| 11 | 18 | 330.9981909 | 11157.84298 | 1 |
| 12 | 19 | 329.153717 | 9677.357379 | 0.994618307 |
| 13 | 22 | 324.9606133 | 8625.344351 | 1 |
| 14 | 24 | 337.5837533 | 10591.97258 | 0.987143291 |
| 15 | 27 | 338.2493291 | 11318.75341 | 0.545666493 |
| 16 | 28 | 344.6051413 | 18976.94823 | 0.833337083 |
| 17 | 29 | 338.8286353 | 12191.84808 | 0.52245813 |
| 18 | 31 | 337.0006355 | 10285.08031 | 0.525828839 |
| 19 | 33 | 333.6776173 | 10298.6094 | 0.815941885 |
| 20 | 34 | 331.1310386 | 8244.80057 | 0.694781646 |
| 21 | 35 | 335.2997715 | 8809.206962 | 0.566857963 |

$$y(T) = \frac{x_3}{1 + \exp(x_2(\frac{1}{T+273} - \frac{1}{x_1}))} + (1 - x_3),$$

| # | Thermometer # | $x_1$ | $x_2$ | $x_3$ |
|---|---|---|---|---|
| 22 | 36 | 336.1214468 | 11624.57974 | 0.836516298 |
| 23 | 37 | 353.3445966 | 18871.30428 | 0.830113314 |
| 24 | 39 | 338.6427844 | 12726.85632 | 0.688291456 |
| 25 | 40 | 349.1060106 | 15833.30373 | 0.820629323 |
| 26 | 41 | 335.0478148 | 9560.32221 | 0.672643612 |
| 27 | 42 | 339.6166315 | 11086.98181 | 0.634399558 |
| 28 | 43 | 342.8479158 | 13291.4851 | 0.634225601 |
| 29 | 44 | 337.0541991 | 10000.54106 | 0.694045146 |
| 30 | 46 | 336.8142905 | 11999.34187 | 0.603483646 |
| 31 | 47 | 343.4674461 | 12224.56877 | 0.62093385 |
| 32 | 48 | 342.212854 | 15465.90574 | 0.586838051 |
| 33 | 50 | 335.324373 | 10402.40603 | 0.64782752 |
| 34 | 51 | 335.8328327 | 10633.55758 | 0.67955356 |
| 35 | 52 | 336.1428879 | 11288.18179 | 0.65561394 |
| 36 | 53 | 342.7548762 | 11779.99274 | 0.83415655 |
| 37 | 55 | 341.3698428 | 12456.20342 | 0.643154134 |
| 38 | 56 | 344.1170518 | 11984.37818 | 0.829504282 |
| 39 | 57 | 338.2690616 | 11548.34826 | 0.664227789 |
| 40 | 59 | 333.6785009 | 7985.542733 | 0.916788066 |
| 41 | 61 | 329.8452601 | 8235.799671 | 0.797849971 |
| 42 | 62 | 330.7985286 | 8988.639691 | 0.805636575 |

$$y(T) = \frac{x_3}{1 + \exp(x_2(\frac{1}{T+273} - \frac{1}{x_1}))} + (1 - x_3),$$

| # | Thermometer # | $x_1$ | $x_2$ | $x_3$ |
|---|---|---|---|---|
| 43 | 64 | 330.5539297 | 8731.492906 | 0.709344298 |
| 44 | 73 | 329.8168418 | 9322.997341 | 0.832801816 |
| 45 | 76 | 326.6647068 | 8654.988618 | 0.896300155 |
| 46 | 77 | 329.4865995 | 9675.248056 | 0.820042664 |
| 47 | 79 | 331.9733374 | 10999.89849 | 0.678104607 |
| 48 | 81 | 328.9760406 | 9526.357972 | 0.823458076 |
| 49 | 84 | 328.5942553 | 9580.136644 | 0.829931216 |
| 50 | 87 | 349.4463917 | 19647.18768 | 0.667374118 |
| 51 | 100 | 325.1423503 | 7585.871382 | 0.99316774 |
| 52 | 102 | 339.3814551 | 10689.31137 | 0.990852592 |
| 53 | 109 | 336.4736748 | 10650.38749 | 0.728366931 |
| 54 | 114 | 329.6405164 | 8266.090407 | 0.864031729 |
| 55 | 116 | 327.1720694 | 7579.079528 | 0.941506805 |
| 56 | 135 | 333.5593291 | 9502.132229 | 0.759973354 |

$$y(T) = \frac{x_3}{1 + \exp(x_2(\frac{1}{T+273} - \frac{1}{x_1}))} + \frac{x_6}{1 + \exp(x_5(\frac{1}{T+273} - \frac{1}{x_4}))} + (1 - x_3 - x_6),$$

| # | Thermometer # | $x_1$ | $x_2$ | $x_3$ | $x_4$ | $x_5$ | $x_6$ |
|---|---|---|---|---|---|---|---|
| 1 | 9 | 308.9880083 | 17813.53313 | 0.210083657 | 338.2086081 | 11400.42223 | 0.621085225 |
| 2 | 10 | 281.6673324 | 14020.31762 | 0.146457875 | 339.8420536 | 13690.35584 | 0.815771894 |
| 3 | 12 | 314.8464773 | 14833.1645 | 0.316948568 | 336.63098 | 10878.75221 | 0.676628962 |
| 4 | 14 | 303.3582187 | 11519.79536 | 0.388086055 | 337.2073288 | 10767.55088 | 0.602147731 |
| 5 | 15 | 302.8896532 | 12230.188 | 0.397484566 | 337.6399776 | 10773.46712 | 0.598794083 |
| 6 | 16 | 303.5960164 | 12357.23069 | 0.398196206 | 337.2506722 | 10716.7563 | 0.59910514 |
| 7 | 17 | 306.3559507 | 17208.01508 | 0.238727786 | 338.2199992 | 11741.41786 | 0.650831297 |
| 8 | 20 | 309.4257758 | 16236.0265 | 0.338687296 | 337.4795938 | 11314.40049 | 0.65857739 |
| 9 | 21 | 306.3559507 | 17208.01508 | 0.238727786 | 338.2199992 | 11741.41786 | 0.650831297 |
| 10 | 23 | 313.1278483 | 6325.606669 | 0.745470717 | 342.43388 | 17123.8163 | 0.197494514 |
| 11 | 25 | 306.3559507 | 17208.01508 | 0.238727786 | 338.2199992 | 11741.41786 | 0.650831297 |
| 12 | 26 | 289.2273495 | 9949.146249 | 0.132866393 | 337.2175076 | 11973.07884 | 0.61000224 |
| 13 | 30 | 289.0625866 | 9847.903295 | 0.132807619 | 337.6748922 | 10854.41476 | 0.480890528 |
| 14 | 32 | 273.0000043 | 5000 | 0.010000003 | 331.6233731 | 8119.144141 | 0.504927011 |
| 15 | 38 | 297.7650002 | 13043.52222 | 0.201192067 | 346.7915028 | 14734.26603 | 0.706624382 |
| 16 | 45 | 302.4885264 | 15133.8508 | 0.167457391 | 337.9352318 | 11745.75065 | 0.591731958 |
| 17 | 49 | 302.4885264 | 15133.8508 | 0.167457391 | 337.9352318 | 11745.75065 | 0.591731958 |
| 18 | 54 | 305.0073647 | 11662.70993 | 0.261890146 | 343.1943502 | 12816.82972 | 0.735801766 |
| 19 | 58 | 295.4535426 | 8965.180951 | 0.280815185 | 337.924395 | 12149.01657 | 0.620541073 |
| 20 | 60 | 293.2110177 | 9516.991648 | 0.260099587 | 339.1295384 | 12159.96733 | 0.64810612 |
| 21 | 63 | 300.3605963 | 13265.06061 | 0.245190831 | 336.37097 | 10714.92635 | 0.670897388 |

$$y(T) = \frac{x_3}{1+\exp(x_2(\frac{1}{T+273} - \frac{1}{x_1}))} + \frac{x_6}{1+\exp(x_5(\frac{1}{T+273} - \frac{1}{x_4}))} + (1 - x_3 - x_6),$$

| # | Thermometer # | $x_1$ | $x_2$ | $x_3$ | $x_4$ | $x_5$ | $x_6$ |
|---|---|---|---|---|---|---|---|
| 22 | 65 | 311.9589384 | 19898.86956 | 0.213034765 | 344.8254578 | 11560.82381 | 0.622035324 |
| 23 | 66 | 300.5940171 | 11691.08639 | 0.383444497 | 349.4863889 | 11036.42305 | 0.61891437 |
| 24 | 67 | 315.4273724 | 15639.91098 | 0.381330147 | 341.6736752 | 12355.26075 | 0.616796709 |
| 25 | 68 | 296.3668116 | 11347.32563 | 0.39828988 | 350.0251325 | 8242.904675 | 0.592288195 |
| 26 | 69 | 302.3314354 | 13818.18536 | 0.237981054 | 344.2037247 | 12360.38261 | 0.593574319 |
| 27 | 70 | 300.7375431 | 13341.22923 | 0.365563081 | 338.684499 | 12058.69023 | 0.626644816 |
| 28 | 71 | 286.5877484 | 12885.83652 | 0.300734316 | 345.062486 | 14310.68094 | 0.675980331 |
| 29 | 72 | 302.1379939 | 12458.71999 | 0.364459728 | 355.3088846 | 9314.205183 | 0.626685664 |
| 30 | 74 | 299.7908184 | 15722.48654 | 0.19579424 | 331.7359107 | 10838.67035 | 0.79991197 |
| 31 | 75 | 302.3828335 | 13486.76948 | 0.448749068 | 350.1560366 | 11509.89288 | 0.547568948 |
| 32 | 78 | 299.5400662 | 12767.28614 | 0.26776744 | 332.6599457 | 12282.22718 | 0.721346361 |
| 33 | 80 | 298.5534084 | 14080.11332 | 0.211871235 | 332.9196432 | 11636.90487 | 0.778114394 |
| 34 | 82 | 297.6216439 | 11680.06062 | 0.274423312 | 332.974836 | 12577.58988 | 0.70451426 |
| 35 | 83 | 298.8406257 | 16853.55431 | 0.510655389 | 341.9519339 | 13096.78272 | 0.491703422 |
| 36 | 85 | 284.7991565 | 9071.954336 | 0.185110908 | 346.2284315 | 16660.3831 | 0.649545771 |
| 37 | 86 | 300.7344977 | 12834.53835 | 0.300711823 | 334.5027085 | 11890.61447 | 0.690801422 |
| 38 | 88 | 294.0920261 | 12984.28247 | 0.330897182 | 343.6241675 | 13400.00931 | 0.663940139 |
| 39 | 89 | 307.5844579 | 10906.15722 | 0.331809489 | 348.5005776 | 10886.55314 | 0.505382813 |
| 40 | 90 | 284.0626211 | 5004.51419 | 0.236956986 | 335.9782674 | 11186.8655 | 0.66248094 |
| 41 | 91 | 298.2019576 | 8659.450164 | 0.409389752 | 342.9375674 | 12397.92092 | 0.576231516 |
| 42 | 92 | 292.06445 | 9927.611337 | 0.264333748 | 336.3371551 | 11359.94801 | 0.723334104 |

$$y(T) = \frac{x_3}{1+\exp(x_2(\frac{1}{T+273} - \frac{1}{x_1}))} + \frac{x_6}{1+\exp(x_5(\frac{1}{T+273} - \frac{1}{x_4}))} + (1 - x_3 - x_6),$$

| # | Thermometer # | $x_1$ | $x_2$ | $x_3$ | $x_4$ | $x_5$ | $x_6$ |
|---|---|---|---|---|---|---|---|
| 43 | 93 | 301.9251021 | 13161.54256 | 0.255451663 | 347.6784256 | 11046.40164 | 0.568420175 |
| 44 | 94 | 295.0413898 | 10455.09093 | 0.276648999 | 338.7533491 | 11208.14784 | 0.648404188 |
| 45 | 95 | 291.1005171 | 7417.715041 | 0.422964677 | 344.419995 | 11638.21137 | 0.562137412 |
| 46 | 97 | 299.0944752 | 9793.243765 | 0.250685995 | 347.3101391 | 10917.26934 | 0.563625378 |
| 47 | 98 | 292.2321057 | 10600.90497 | 0.201939802 | 335.4102022 | 9334.410221 | 0.629500553 |
| 48 | 99 | 290.5141322 | 9505.387115 | 0.339739807 | 338.415424 | 11108.35215 | 0.553193491 |
| 49 | 101 | 292.0639496 | 10236.80481 | 0.380685319 | 338.2003051 | 10981.85002 | 0.567990121 |
| 50 | 103 | 290.4038583 | 7455.540707 | 0.248877808 | 342.5012902 | 12947.30391 | 0.614342998 |
| 51 | 104 | 299.9031775 | 14027.92833 | 0.357504492 | 340.5208558 | 11502.26857 | 0.622026404 |
| 52 | 105 | 302.1303604 | 15582.99299 | 0.348972809 | 337.7497955 | 11153.816 | 0.647513336 |
| 53 | 106 | 282.3075523 | 8147.145555 | 0.46104691 | 343.2536682 | 12111.00826 | 0.497268086 |
| 54 | 107 | 319.2365328 | 16455.71151 | 0.270787833 | 340.6369144 | 12312.53398 | 0.725711415 |
| 55 | 108 | 304.8670056 | 13884.35792 | 0.382802755 | 338.8713371 | 11907.61908 | 0.608558516 |
| 56 | 110 | 294.9662089 | 14098.43484 | 0.383782391 | 336.6268466 | 10916.30721 | 0.601951006 |
| 57 | 111 | 299.1293394 | 10263.26189 | 0.49073484 | 346.1561242 | 12100.36556 | 0.471260015 |
| 58 | 112 | 298.6089342 | 11102.68555 | 0.376234811 | 340.2169667 | 10310.39469 | 0.604865862 |
| 59 | 113 | 299.2410264 | 9317.489799 | 0.368663229 | 338.5067542 | 11896.37451 | 0.613182826 |
| 60 | 115 | 313.6542954 | 16862.26222 | 0.620169313 | 344.839492 | 12577.05701 | 0.373926919 |
| 61 | 117 | 343.5636821 | 12003.34883 | 0.538539111 | 320.0918252 | 16184.55517 | 0.457457851 |
| 62 | 118 | 343.3078027 | 11898.49472 | 0.541601037 | 319.983094 | 16262.51865 | 0.454429502 |
| 63 | 119 | 303.4049259 | 11929.71261 | 0.493461138 | 340.3643222 | 10759.84119 | 0.496310682 |

$$y(T) = \frac{x_3}{1+\exp(x_2(\frac{1}{T+273} - \frac{1}{x_1}))} + \frac{x_6}{1+\exp(x_5(\frac{1}{T+273} - \frac{1}{x_4}))} + (1 - x_3 - x_6),$$

| # | Thermometer # | $x_1$ | $x_2$ | $x_3$ | $x_4$ | $x_5$ | $x_6$ |
|---|---|---|---|---|---|---|---|
| 64 | 120 | 310.1916145 | 14304.25191 | 0.388890072 | 344.21569 | 12348.05446 | 0.603438854 |
| 65 | 121 | 305.4829954 | 16642.97135 | 0.325764236 | 337.2034161 | 11181.08537 | 0.669788973 |
| 66 | 122 | 299.7526133 | 11823.74 | 0.298048289 | 337.1835336 | 11450.92326 | 0.660910419 |
| 67 | 123 | 306.1567239 | 13862.9129 | 0.351371319 | 339.0028898 | 11153.90613 | 0.599308972 |
| 68 | 124 | 312.61455 | 17314.75371 | 0.453452757 | 347.6625668 | 13990.34847 | 0.540745021 |
| 69 | 125 | 305.4829954 | 16642.97135 | 0.325764236 | 337.2034161 | 11181.08537 | 0.669788973 |
| 70 | 126 | 284.6510691 | 5000.045195 | 0.403575239 | 336.0911385 | 11492.55708 | 0.637172077 |
| 71 | 128 | 286.2055871 | 5000.000057 | 0.347555698 | 338.7661204 | 11467.1249 | 0.616393182 |
| 72 | 129 | 305.4829954 | 16642.97135 | 0.325764236 | 337.2034161 | 11181.08537 | 0.669788973 |
| 73 | 130 | 304.159842 | 15894.33201 | 0.31756254 | 336.7714998 | 11183.71113 | 0.678436591 |
| 74 | 131 | 303.2564014 | 15591.62637 | 0.338811702 | 337.4616291 | 11277.48543 | 0.657098027 |
| 75 | 132 | 300.2015556 | 15106.38053 | 0.340752685 | 339.0402502 | 10892.454 | 0.656873977 |
| 76 | 133 | 300.1846214 | 14484.0367 | 0.360726163 | 338.965506 | 11336.13955 | 0.635318514 |
| 77 | 134 | 297.3962281 | 12359.35301 | 0.366865835 | 340.0286212 | 11375.67491 | 0.630353672 |
| 78 | 136 | 296.9422692 | 13072.73222 | 0.321524667 | 339.9352858 | 11790.42558 | 0.675267533 |
| 79 | 0 | 300.6547657 | 12468.07608 | 0.327284504 | 337.1527045 | 11213.40975 | 0.664692809 |
